# A stochastic oscillator model simulates the entrainment of vertebrate cellular clocks by light

**DOI:** 10.1101/2021.03.24.436598

**Authors:** Vojtěch Kumpošt, Daniela Vallone, Srinivas Babu Gondi, Nicholas S. Foulkes, Ralf Mikut, Lennart Hilbert

## Abstract

The circadian clock is a cellular mechanism that synchronizes various biological processes with respect to the time of the day. While much progress has been made characterizing the molecular mechanisms underlying this clock, it is less clear how external light cues influence the dynamics of the core clock mechanism and thereby entrain it with the light-dark cycle. Zebrafish-derived cell cultures possess clocks that are directly light-entrainable, thus providing an attractive laboratory model for circadian entrainment. Here, we have developed a stochastic oscillator model of the zebrafish circadian clock, which accounts for the core clock negative feedback loop, light input, and the proliferation of single-cell oscillator noise into population-level luminescence recordings. The model accurately predicts the entrainment dynamics observed in bioluminescent clock reporter assays upon exposure to a wide range of lighting conditions. Furthermore, we have applied the model to obtain refitted parameter sets for cell cultures exposed to a variety of pharmacological treatments and predict changes in single-cell oscillator parameters. Our work paves the way for model-based, large-scale screens for genetic or pharmacologically-induced modifications to the entrainment of circadian clock function.

**Author summary:** The circadian clock is a key, cell-autonomous timing mechanism that is encountered in most organisms. It is entrained by environmental lighting conditions and in turn temporally coordinates most aspects of physiology according to the time of day. Cell lines derived from zebrafish are attractive experimental models for studying how clocks are entrained by light since they possess clocks that respond directly to light stimuli. Here we describe a mathematical model for the behavior of the circadian clock in zebrafish cell lines during exposure to a range of lighting conditions. Using this model, we can determine how different pharmacological treatments may affect the entrainment dynamics of the clock and the degree of synchronization of individual cells’ circadian clocks in bioluminescent clock reporter assays. Our current model is mathematically simple and thus easy to apply and extend in future studies.

## Introduction

The circadian clock is a timekeeping mechanism that temporally coordinates the majority of biological processes according to the environmental day-night cycle [1]. At the molecular level, the core of this mechanism is based on a set of clock genes and proteins which constitute an autoregulatory, transcription-translation negative feedback loop [2]. In vertebrates, the positive elements Clock and Bmal are transcriptional activators that heterodimerize, bind to E-box enhancers, and thereby activate the transcription of a set of negative regulatory genes, which encode cryptochrome (Cry) and period (Per) proteins. Cry and Per directly interact and interfere with the Clock-Bmal activators, in consequence blocking transcriptional activation. This results in a reduction in *cry* and *per* gene expression, lowering of the nuclear levels of Per and Cry, leading ultimately to the release of inhibition of the Clock-Bmal heterodimers. This feedback cycle takes approximately 24 hours to complete until it can start again, in the process generating an autonomous 24-hour rhythm. Since the period of this rhythm is not precisely 24 hours, it relies upon daily resetting to ensure synchronization with the environmental day-night cycle. Environmental signals that are indicative of the time of day (zeitgebers) are transduced to the core clock mechanism via various signaling pathways and thereby entrain the clock [3]. Light represents the primary zeitgeber in most organisms and accordingly has received significant attention [4]. In mammals, light is perceived by a dedicated circadian photoreceptor, the non-visual opsin melanopsin which is expressed in intrinsically photosensitive retinal ganglion cells (ipRGCs) in the retina [5]. Information on external light levels is then relayed indirectly via the retinohypothalamic tract (RHT) to the central clock located in the suprachiasmatic nucleus (SCN) within the hypothalamus, resulting in induced expression of the *per1* and *per2* clock genes, which in turn sets the phase of the SCN clock. The adjusted phase of the SCN clock is then communicated to other peripheral tissue clocks by a complex combination of systemic signals [6]. Frequent disruption of circadian clock function by exposure to irregular light-dark (LD) cycles can lead to mood and cognitive disorders [7] as well as a broad range of human pathologies including metabolic disorders and cancer [8].

Mathematical modeling has provided much insight into the regulatory mechanisms underlying the circadian clock, including the effects of pharmacological treatments and light entrainment [9]. Even though detailed mathematical models of the molecular mechanisms underlying circadian rhythms exist [10–13], it is often favorable to use simpler models that condense the underlying mechanism into a few effective elements that are sufficient to reproduce experimentally-observed behavior [14]. Such minimal models have previously been employed to study various aspects of the circadian clock, including light entrainment [15], the effect of noise [16], and the coupling of cellular feedback loops [17]. Intrinsic cellular noise leads to stochastic behavior of the circadian clock [18]. Even in bioluminescent reporter assays that record average signals from large cell populations, the noise in single-cell dynamics has a significant impact on clock rhythmicity when cells are placed in constant darkness (free-running conditions) [19]. In the absence of synchronizing zeitgebers, the single-cell oscillators progressively desynchronize, leading to the population-level loss of amplitude, even though individual cells maintain oscillation with undiminished amplitude [20]. Re-exposure to a zeitgeber has the effect of resynchronizing the asynchronous single-cell oscillators with the appropriate phase and rhythmicity re-emerges at the population level. Experimental measurement of the degree of single-cell clock synchronization requires long-term imaging of individual cells, which is technically challenging and not suited to large-scale screening experiments. Here, a mathematical modeling approach that could predict the degree of individual cell clock synchronization from population-level recordings would be valuable.

Fish represent attractive model organisms for studying circadian clocks in vertebrates. They constitute the largest and most diverse vertebrate group with species living under a wide range of environmental conditions. Therefore, they enable the study of how clocks have adapted to changing environmental conditions over the course of evolution [21]. Furthermore, a better understanding of circadian clock regulation and function in fish has potential benefits for aquaculture where lighting conditions have long been recognized to have a major impact on the rate of growth [22]. In addition, zebrafish is a particularly attractive genetic model organism to study the molecular basis of how the circadian clock is regulated by light. Specifically, zebrafish tissues and even cell lines possess directly light-entrainable clocks [23]. Unlike mammalian cells which require transient pharmacological treatments to synchronize cell culture clocks [24, 25], in the case of zebrafish cells, the clocks can be regulated non-invasively simply by changing lighting conditions. These light-responsive cell lines are also suitable for high-throughput screening as well as studies of the transcriptional control mechanisms mediating light entrainment [26]. In this regard, many bioluminescent reporter systems have now been established in zebrafish cell lines and enable the non-invasive assessment of dynamic changes in clock gene transcription at high temporal resolution over the course of light exposure protocols [27]. A previous theoretical model of the zebrafish circadian clock has shown that, in principle, simulated single-cell oscillations can be used to reconstruct average luminescence signals from cell populations [28]. However, this model is based on the consideration of two interlocked feedback loops and a large number of parameters, making reliable fitting and readjustment of model parameters to limited experimental data difficult.

Here, we present a minimal stochastic oscillator model of the circadian clock in zebrafish cell lines. This model, based on the classic Goodwin model [29], contains only three variables but reproduces and predicts the main oscillatory characteristics observed in bioluminescent reporter assays under a broad range of experimental lighting conditions. We further adjusted the model parameters to infer the impact of various pharmacological treatments on core clock dynamics at the cellular level. This approach paves the way towards model-based, large-scale screens for genetically or pharmacologically-induced modifications affecting the degree of synchronization of single-cell circadian oscillators.

## Results

### Characterization of light entrainment in cell culture

To explore the entrainment dynamics of the vertebrate circadian clock in an experimental model, we used zebrafish PAC-2 cell lines. PAC-2 cellular clocks can be entrained by direct exposure to light and they have been widely used to study the transcriptional mechanisms of light entrainment [26, 30, 31]. We chose to study the dynamics of circadian clock gene expression by non-invasive luminescence assays of PAC-2 cells stably transfected with a *zper1b* promoter bioluminescent reporter construct [26]. In these cells, reporter gene transcription is directly controlled by the core clock mechanism via E-box enhancers. To obtain a data set that supports the development, adjustment, and validation of a mathematical model, we recorded luminescence over several days, in cells exposed to a variety of different lighting conditions (Fig 1A and S1 Fig), including various LD cycles, constant darkness, and constant light. The resulting data set contains an extensive number of repeats (S1 Table), further supporting model fitting and validation.

**Fig 1.**
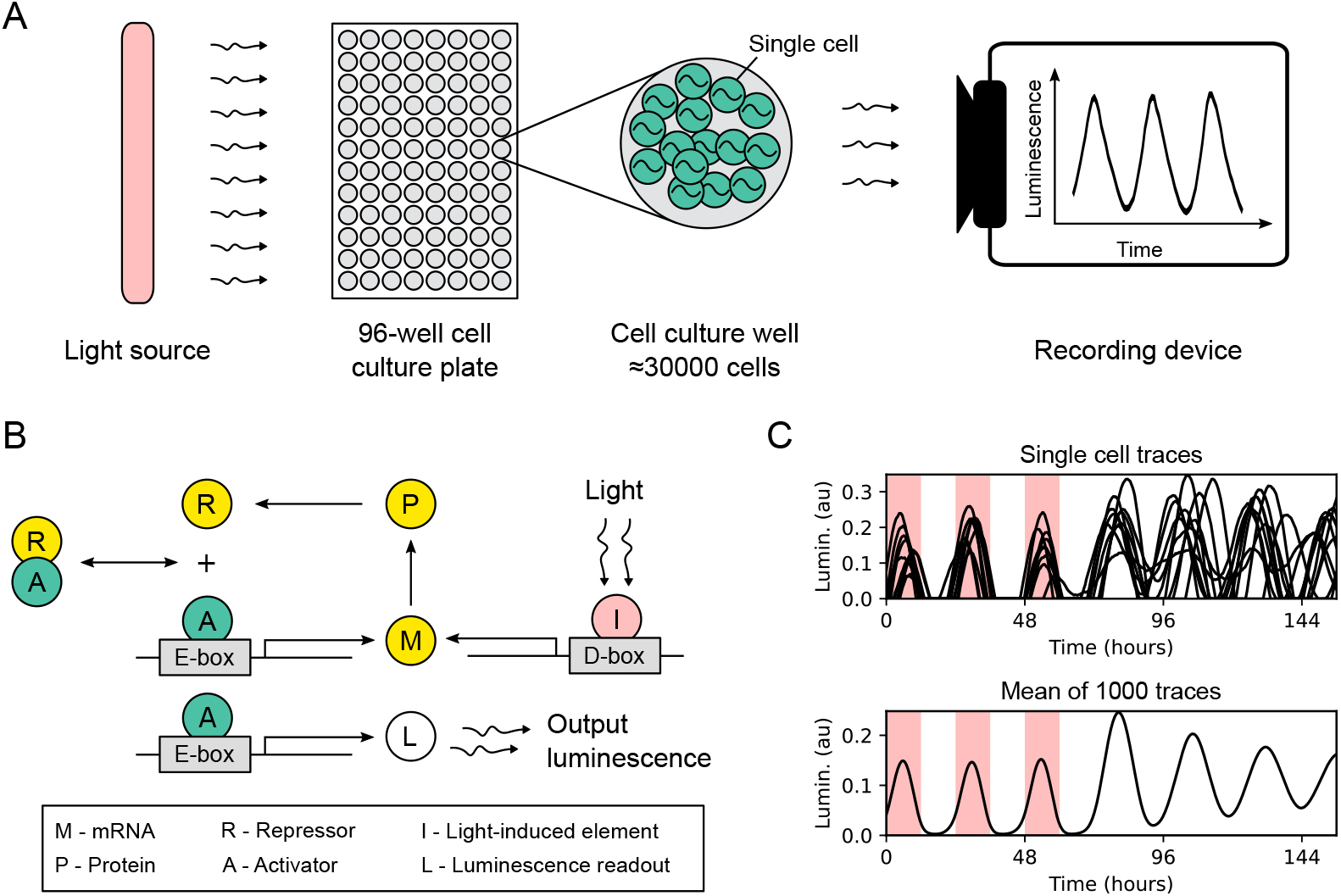
Experiment setup and simulation of core circadian clock dynamics in zebrafish cell cultures. (A) 96-well culture plates were placed in a dark room and illuminated with a time-controlled light source. Each well contains approximately 30000 cells transfected with a bioluminescent reporter of *zper1b* transcription. The output luminescence from each well is recorded as a separate time trace. (B) Schematic of the mathematical model of the zebrafish core circadian clock. The activator (green) binds to the E-box enhancer in the promoter of a clock gene and activates the production of the repressor (yellow). After transcription, translation, and translocation back to the nucleus, the repressor binds to the activator, thus preventing E-box-driven transcriptional activation. External light stimuli take effect through the activation of a light-driven gene with the D-box enhancer. The luminescence output is assumed to be proportional to the E-box activation. (C) Simulated luminescence traces are obtained by averaging 1000 independent evaluations of the model. The output luminescence after normalization is presented in arbitrary units (au). Red shading indicates periods when the light source was turned on.

### Stochastic oscillator model

As a conceptual basis for the development of a mathematical model for the PAC-2 cell clock, we considered that the transcription-translation negative feedback loop at the core of the zebrafish circadian clock consists of positive elements (activators) and negative elements (repressors). The activators bind to E-box enhancers located in the promoter region of clock genes, thus inducing the transcription of the repressors. After transcription, translation, and translocation back to the nucleus, the repressors physically interact with the activators and thereby inhibit the transcriptional induction of their own genes [32]. As a first step, we describe the core clock mathematically using three coupled differential equations connected in a negative feedback loop (Fig 1B, Eq 1). This type of model is known as the Goodwin oscillator [29] and is frequently used as an abstraction of the core clock [33]. The three state variables implicitly encode the delay that is necessary to produce the oscillatory behavior [34]. To more closely connect our mathematical model to the known regulatory processes, we arbitrarily label the three variables as mRNA (*M*), cytoplasmic protein (*P*), and nuclear protein that acts as a repressor (*R*). The repressor *R* binds to the activator *A*, forming an inactive *RA* complex with 1:1 stoichiometry [13], thus preventing the binding of the activator to the E-box enhancers, resulting in inhibition of gene expression. We use an according protein sequestration function (f, Eq 2) to describe the transcriptional activation [35]. This function defines the transcriptional activation as a function of the repressor and activator concentrations. The light input is implemented as a light-induced element *I*, whose activation increases the production of mRNA M. Similar extensions to the Goodwin model are commonly used to represent light stimuli [15,36,37]. Specifically, in our model, *I* represents light-driven activation of a subset of *per* and *cry* clock genes that is characteristic of the zebrafish circadian clock and is mediated by D-box enhancers [30,31]. As a result of this gene activation, there is an increase in the production of negative elements. The function f is the quantity in our model that most closely represents the induction of the *zper1b:luc* reporter by E-box enhancers. We thus use the value of the function f to represent the luminescence read-out obtained in our experiments.

Each well of a 96-well culture plate contains approximately 30000 independently oscillating cells that contribute to the luminescence signal measured from that well. Intrinsic cellular noise affects the individual oscillators but is averaged in the population-level luminescence recordings [20]. One classical phenomenon demonstrating the relevance of single-cell dynamics is the population-level progressive loss of the amplitude of oscillations in constant darkness while individual cells continue to oscillate with undiminished amplitude (Fig 1C). We have therefore implemented noise at the single-cell level by additive noise terms in our model [16]. To mimic the averaging over a population of single-cell oscillators in the luminescence assays, we simulate 1000 instances of our model and obtain the mean value as the final output [19,28].

### Parameter adjustment and model validation

We next adjusted the model parameters to luminescence recordings with a custom fitting algorithm based on an evolutionary optimizer. Specifically, we compared the behavior of our model with luminescence recordings showing three characteristic phenomena: attenuation of the oscillation amplitude during free-running conditions in constant darkness, synchronization of the phase of the rhythm upon transfer from constant darkness to an LD cycle, and oscillation repression under constant light. We first adjusted our model parameters to luminescence recordings with varied light conditions and evaluated the goodness of fit by the model efficiency coefficient (*E_f_*, Eq (14)). The resulting model efficiency was high (16 wells, mean ± standard deviation, *E_f_* = 0.89 ± 0.03, Fig 2A). Without further readjustment of model parameters, we simulated luminescence traces for experiments with different lighting conditions, resulting in fair model efficiency (4 wells, *E_f_* = 0.63 ± 0.21, Fig 2B). Upon visual inspection, the model captured all key behaviors observed in *zper1b:luc* assays. We also tested the performance of the model under LD cycles with period lengths significantly longer and shorter than 24 hours (30 hours, 15:15 LD; and 20 hours, 10:10 LD). In the case of our model, the goodness of fit was lower than under the 12:12 LD cycle (32 wells, 15:15 LD, *E_f_* = 0.24 ± 0.20; 16 wells, 10:10 LD, *E_f_* = 0.60 ± 0.11; Fig 2C), mainly due to mismatched profiles of each expression cycle. Nevertheless, the model was still entrained and oscillated with the corresponding period of these long and short LD cycles. This is consistent with our previous experiments where we showed that the PAC-2 cell clock exhibits rhythmicity under these conditions with an adjustment of the period length of the reporter rhythm to match the period of the LD cycle [26]. Therefore, the model can reproduce and predict characteristics of the entrainment of the endogenous PAC-2 cellular clock under complex lighting conditions and loses its accuracy only for entraining period lengths that differ significantly from the natural 12:12 LD cycle.

**Fig 2.**
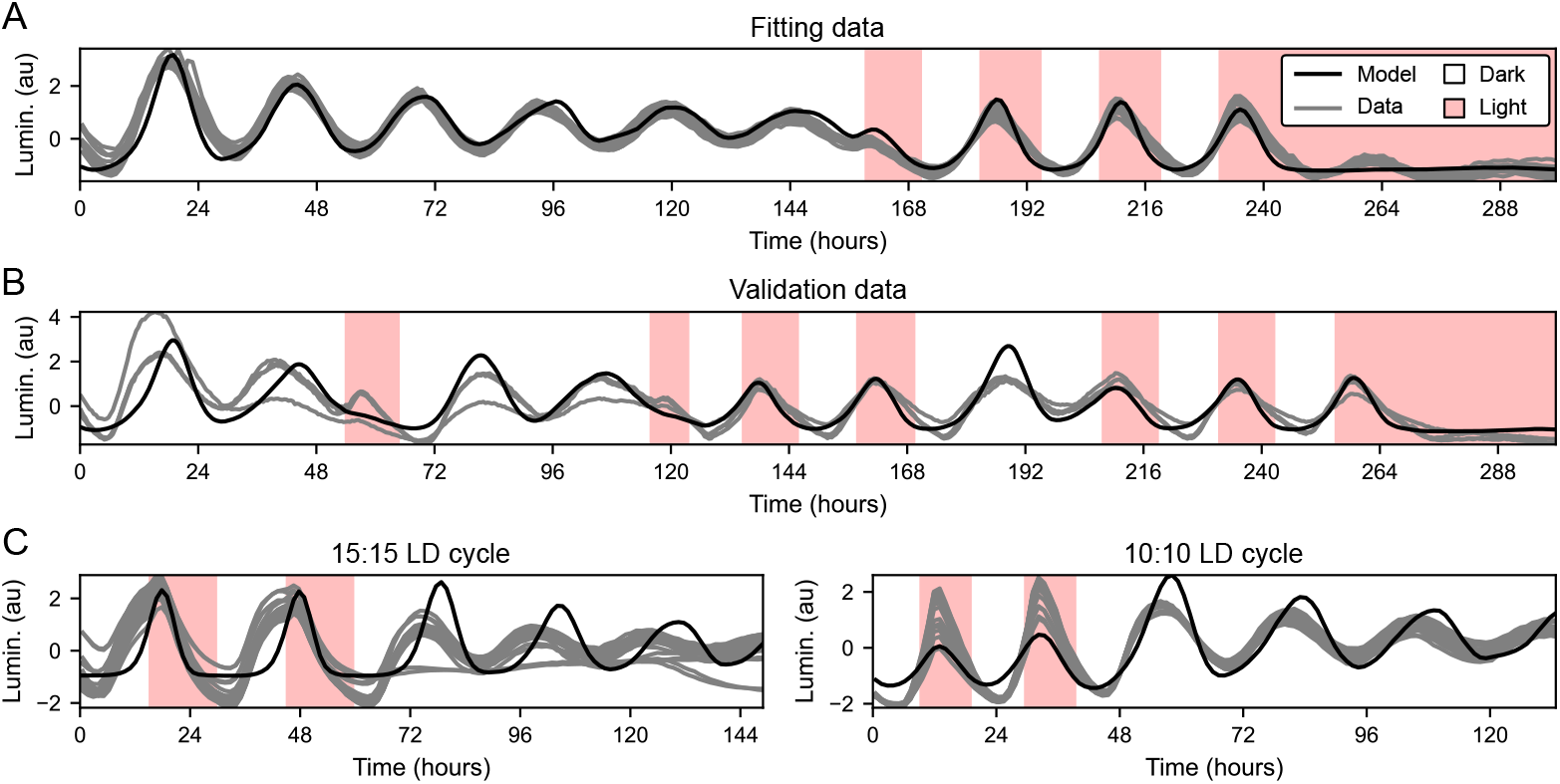
Simulation of *zper1b:luc* expression dynamics under various lighting conditions. (A) On this plate, cell cultures were exposed to six days of constant darkness, three 12:12 LD cycles, and three days of constant light. This data was used to estimate the model parameters. (B) On this plate, cell cultures were exposed to a combination of LD cycles, constant darkness, and constant light. The same parameters as in panel A were used to run the model, showing that the model can also reproduce dynamics of data not used for parameter fitting. (C) These plates were exposed to 15:15 and 10:10 LD cycles. Again, parameters estimated from panel A were used without further fitting.

### Representing effects of pharmacological treatments as model parameter changes

We next wished to explore whether our mathematical model could discriminate between changes in basic properties of the clock mechanism induced by different pharmacological treatments. We treated our clock reporter PAC-2 cells with six compounds that affect three major signaling pathways regulating the core clock. Specifically, we used forskolin (FOR) and dibutyryl cAMP (DBC) as activators of the cAMP pathway; epidermal growth factor (EGF) and U0126 as an activator and an inhibitor of the MAPK pathway, respectively, as well as phorbol-12-myristate-13-acetate (PMA) and ro-318220 (RO) as an activator and an inhibitor of the PKC pathway, respectively. All these pathways have been previously implicated in light entrainment [38–41]. We assessed the effect of the exposure to these compounds under two different sets of lighting conditions (Fig 3A, B). Starting with the model parameters from our initial adjustment to data obtained from untreated cells, we executed a step-wise refitting to the results after pharmacological treatments. Each compound was administrated at three different concentrations (S1 Table), and parameters for the individual compound concentrations were estimated iteratively starting with the lowest compound concentrations and finishing with the highest compound concentrations. We set the threshold for a good fit as *E_f_* > 0, which excluded 4 of 20 treatments from our further analysis. The accepted and excluded fits are listed in S2 Table. The final model fits for all pharmacological treatments are depicted in S3 Fig.

**Fig 3.**
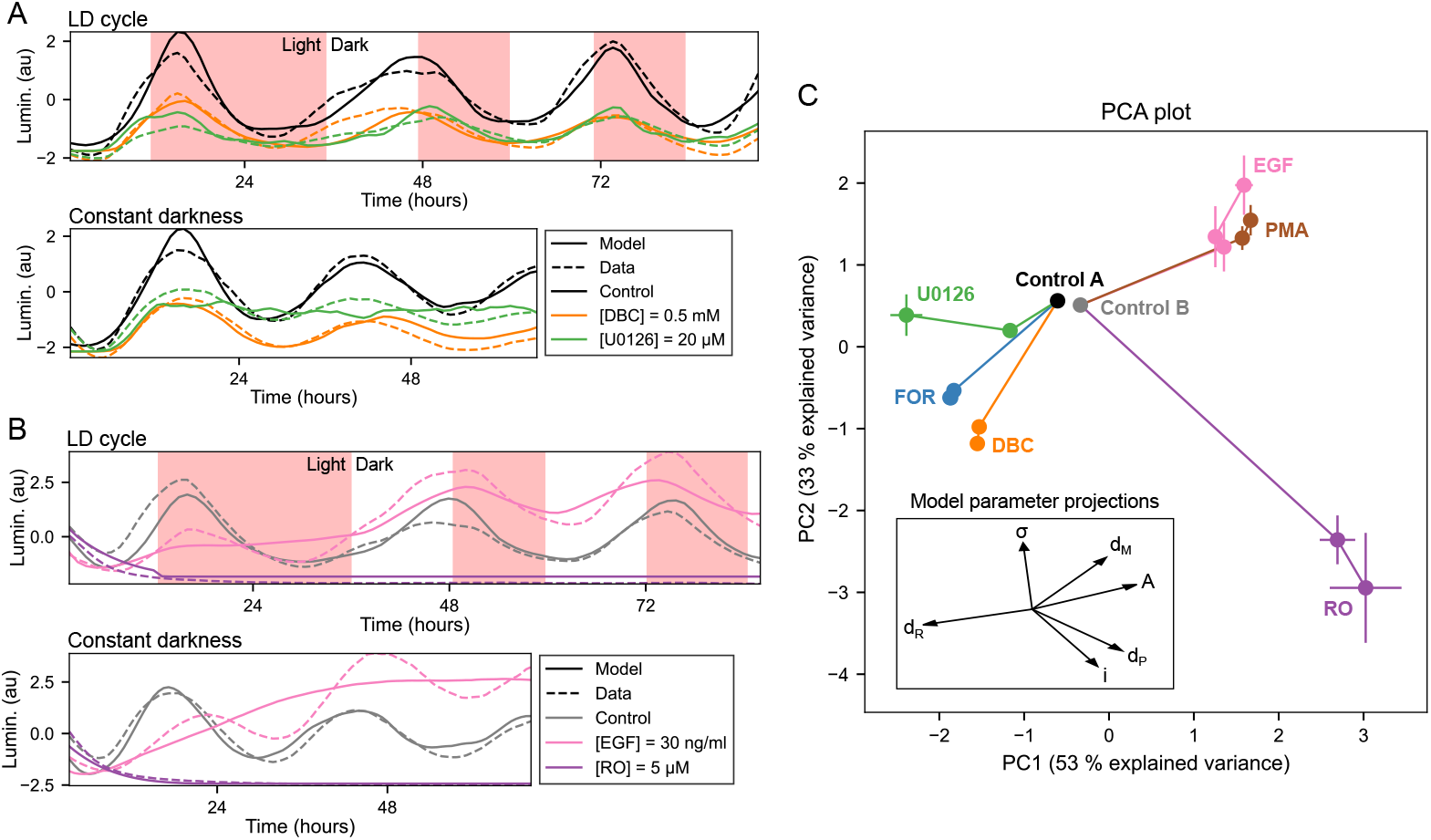
Characterization of compound effects by refitting of model parameters. (A) Representative experimental data traces and model fits for plate set A. Plate set A contained treatments with dimethyl sulfoxide (DMSO) control, forskolin (FOR), dibutyryl cAMP (DBC), and U0126. See S3 FigA, B, C for model fits to all concentrations of FOR, DBC, and U0126. (B) Representative experimental data traces and model fits for plate set B. Plate set B contained treatments with DMSO control, epidermal growth factor (EGF), phorbol-12-myristate-13-acetate (PMA), and ro-318220 (RO). See S3 FigD, E, F for model fits to all concentrations of EGF, PMA, and RO. (C) The principal component plot depicts the projection of the compound-dependent changes in model parameters on two main principal components (PCs), which provides a tool to visualize the changes of the model parameters in a two-dimensional plot. Each compound and concentration is represented with a point. The error bars at each point indicate the spread of the final population of parameter sets obtained by a differential evolution algorithm (median ± median absolute deviation). Individual values used to calculate the median points are shown in S2 Fig. Lines connect increasing concentrations of the same compound, starting from the control condition. Control A is a DMSO control for FOR, DBC, and U0126. Control B is a DMSO control for PMA, EGF, and RO. The explained variance for the first three PCs was 53 %, 33 %, and 8 %.

The principal component analysis (PCA) of the parameter sets representing the individual compounds revealed that the two main principal components together account for 86 % of the explained variance. We can therefore depict the relative changes of the six free parameters in the two-dimensional space defined by the two main principal components (Fig 3C). In the PCA plot, every compound exhibits a change away from the control conditions. Compound-specific changes were increasingly pronounced at higher concentrations, while the overall direction of parameter change was conserved for each of the compounds. Based on the direction of displacement in the principal component plot, different compounds can be grouped. FOR, DBC, and U0126 are all associated with a decrease in principal component 1, reflected in a loss of the oscillation amplitude that is similar for all three compounds (Fig 3A, S3 Fig A, B, C). PMA and EGF are associated with an increase in principal component 1, reflected in a decreased amplitude of the oscillations during the first day of the recording, followed by a sudden transition to a higher amplitude in the subsequent days of the assay (Fig 3B, S3 Fig D, E). RO showed a distinct decrease in principal component 2 and lies considerably further away from all other compounds. This reflects the complete absence of oscillatory behavior associated with this specific treatment (Fig 3B, S3 Fig F). Taken together, our refitting approach allows the categorization of these compounds based on their inferred effects on the model parameters, as indicated by the proximity of compounds with similar effects in the PCA plot.

### Treatment-specific mechanisms of amplitude loss

The impact of the FOR, DBC and U0126 compounds on the rhythmic parameters of the luminescence recordings are similar. However, close inspection of the PCA plot reveals that U0126 lies further away from FOR and DBC (Fig 3C). The change appears to be mainly in the direction of parameter *σ*, suggesting that U0126 differs from FOR and DBC by higher noise intensity. As our model consists of an ensemble of 1000 oscillators, we wondered if the observed difference can be mapped to the altered behavior of the simulated individual cells. As an illustrative example, we focused on comparing the treatments with 0.5 mM DBC and 20 μM U0126. When synchronized cell populations were transferred to constant darkness, both treatments resulted in a more rapid loss of population amplitude than in control-treated cultures. At the level of single cells, however, our model suggests that DBC treatment resulted in reduced single-cell oscillator amplitude, while incubation with U0126 caused a more pronounced desynchronization of individual oscillators due to higher noise (Fig 4). We can thus visualize two distinct sets of single-cell dynamics that both result in the accelerated loss of rhythm amplitude at the population level.

**Fig 4.**
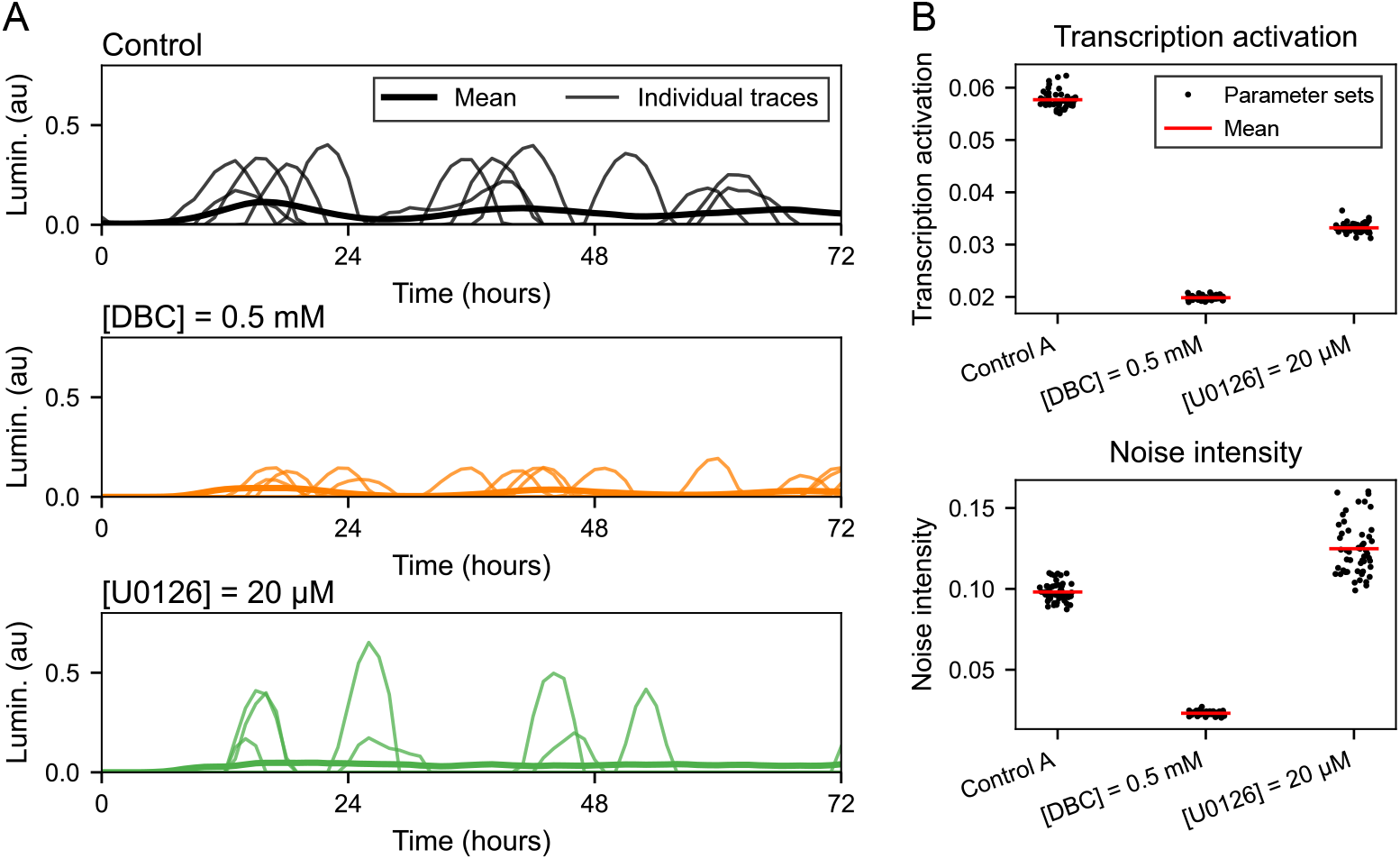
The population-level loss of amplitude is caused by different underlying single-cell mechanisms. (A) Output of the model in constant darkness after previous entrainment by 12:12 LD cycle. For each compound the mean is displayed with 5 of the 1000 individual traces from which the mean was calculated. (B) Transcription activation was calculated as mean of the output luminescence on the depicted time interval. The values for control (*L* = 0.0576 ± 0.0017) are higher than the values for DBC (*L* = 0.0198 ± 0.0004) and U0126 (*L* = 0.0331 ± 0.0012). The noise intensity value for control (*σ* = 0.098 ± 0.006) was lower than for U0126 (*σ* = 0.124 ± 0.016) but higher than for DBC (*σ* = 0.023 ± 0.001). Mean and standard deviation were calculated across the final population of parameter sets obtained by the evolutionary optimizer.

## Discussion

Here, we present a stochastic oscillator model that mimics the dynamic properties of the core circadian clock in cultures of zebrafish cell lines. We find that the model accurately recapitulates the rhythmic properties and entrainment of luminescence recordings under a range of lighting conditions, only losing accuracy when simulating the effects of exposure to LD cycles with periods significantly longer or shorter than 24 hours. Furthermore, we demonstrate the utility of our model for the characterization of luminescence recordings in terms of changes in single-cell core clock regulation.

Our stochastic model is based on the classic three-variable Goodwin model [29] extended by light input and noise terms. We also replaced the Hill function, which is commonly used to describe the negative feedback, with the protein sequestration function, which more closely resembles the molecular mechanisms of transcription inhibition in the vertebrate core clock [13, 35]. The Goodwin model is a minimal abstraction of the core circadian clock and was used in many previous studies to investigate the properties of circadian entrainment [36, 42–44]. Our work shows that this minimal model can simulate the key characteristics of the zebrafish circadian clock, the model simplifies the precise molecular details of clock regulation and only loses predictive strength in experiments where cells are exposed to light cycles with periods that deviate significantly from the natural 12:12 LD cycle. One simplification in our model that is potentially relevant in this respect is the use of only a single feedback loop, even though the actual circadian clock consists of multiple interlocked feedback loops [32]. Furthermore, the core feedback loop consists of multiple clock genes with overlapping functions, which has been predicted to contribute to the regulatory robustness of the clock [45]. However, our results indicate that at a systems level, these extra clock genes effectively act within the context of a complex regulatory circuit that behaves as a single feedback loop.

Cellular noise is an intrinsic part of the clock mechanism [18] and it is essential to consider this factor when interpreting observations made at the cell population level. This importance of cellular noise was also recognized by a previous model of the zebrafish circadian clock [28]. As in our model, signals from an ensemble of stochastic oscillators were averaged to simulate population-level luminescence recordings. We reduced the number of variables (from 5 to 3) and the number of free parameters (from 15 to 6) in relation to the previous model, enabling the application of a fitting algorithm to adjust and quantitatively validate our model against experimental data. The possibility to fit our model to luminescence recordings further provides the foundation for inference of changes in core clock parameters resulting from pharmacological treatments or genetic changes. We used refitting of our model to understand the effect of different compounds on the core clock dynamics. This approach does not identify the specific molecular interactions that are affected by the specific treatments, but rather provides a tool to reveal effective changes in the dynamic properties of the core clock mechanism as an overall regulatory system. This approach is conceptually consistent with previous work suggesting that all light-transduction signaling pathways converge to the same active element [46] and thus exhibit similar effects on the overall clock dynamics.

Inspecting the resulting parameter space, we found that the parameter sets representing the treatments with FOR and DBC were closer to each other than other compounds. This is expected, as both FOR and DBC act as activators of the cAMP pathway and should thus similarly affect the core clock mechanism [47]. The parameter sets representing EGF and PMA were also close to each other. This is consistent with previous work showing that the PKC pathway (activated by PMA) is an upstream activator of the MAPK pathway (activated by EGF) [38, 48]. Those results indicate that our model as well as the fitting procedure can also accurately classify changes in core clock parameters. In addition, our model can predict changes in single-cell dynamics from the population-level luminescence recordings, as seen for the DBC and U0126 treatments. The close proximity of the parameter values for DBC and U0126 lead to similar effects at the population level, namely a reduced amplitude of rhythmic reporter gene expression. However, our model predicts that the effect of U0126 is substantially different from the effect of DBC at the level of the single-cell simulations. While our model predicts that DBC causes the loss of population-level amplitude by inhibition of gene expression at the single-cell level, U0126 is predicted to cause a more rapid desynchronization of the single-cell oscillators.

Assessment of the degree of synchronization of single-cell oscillators in a cell population in culture involves technically challenging and time-consuming imaging that is not trivial to perform as part of a large-scale screen. Within this context, our modeling approach provides the possibility to make rapid predictions about the behavior of individual cell clocks from population-level luminescence recordings. These predictions could then be followed up by more refined imaging-based assays of single-cell dynamics on a much smaller set of samples. Therefore, our mathematical model should contribute a rapid and scalable tool for interpreting the effects of pharmacological treatments and genetic modifications on the circadian clock at the cellular level.

## Materials and methods

### Luminescence recordings

We used PAC-2 light-responsive cells stably transfected with the *zper1b:luc* reporter [26]. 96-well culture plates were seeded with approximately 30000 cells per well and placed in a dark room. Different lighting conditions were applied by exposure to a time-controlled white light source (S1 Fig). Approximately every 40 minutes, each plate was automatically moved into the counting chamber of a TopCount NXT counter (Perkin Elmer) and luminescence was measured for approximately 5 minutes (3 seconds per well). Forskolin (FOR), dibutyryl cAMP (DBC), epidermal growth factor (EGF), U0126, phorbol-12-myristate-13-acetate (PMA) and ro-318220 (RO) were used at concentrations indicated in S1 Table. An overview of the experiment design is shown in Fig 1A.

### Mathematical model

The model equations for a single-cell oscillator represent oscillatory behavior of three clock elements denoted as mRNA *M*, protein *P*, and repressor *R*. The equations read

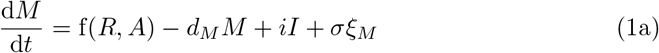

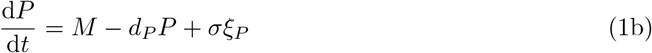

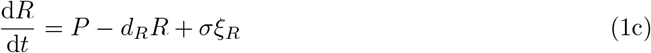

where *A* denotes activator concentration, *d*_*_. are degradation rates, *σ* is noise intensity, and ξ_*_ are independent Wiener processes. *I* is a light input function, which is 0 in darkness and 1 in light. Parameter *i* represents the light sensitivity of the clock. The protein sequestration function f has form [35]

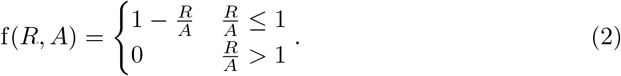

The luminescence produced by a single cell is then equal to

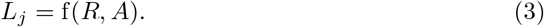

The final model output that corresponds to the luminescence output of a cell culture well is

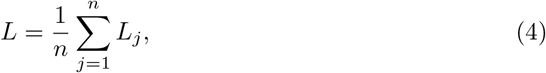

where *j* are independent evaluations of the stochastic model and *n* = 1000.

### Data and model output normalization

Normalization of luminescence recordings is necessary to eliminate variations in amplitude caused by inherent experimental error. We normalized data from untreated cell lines on the trace-by-trace basis by Z-score defined as

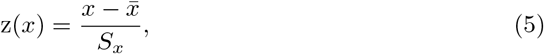

where 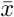 and *S_x_* are sample mean and standard deviation of trace *x*. As the normalized traces of repeats showed very low variance, we used their mean for the model fitting and validation (S4 Fig).

For the plates with compound-treated cell lines, we could not use the trace-by-trace Z-score normalization as the change in mean and variance can be an important aspect of the compound effect. Therefore, we defined the adjusted Z-score

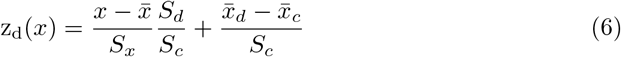

where 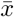 and *S_x_* are sample mean and standard deviation of trace *x*, 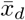 and *S_d_* are mean of means and mean of standard deviations for all traces of the corresponding compound, and 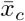 and *S_c_* are mean of means and mean of standard deviations for all control-treated traces. For control the equation reduces to standard Z-score because *S_d_* = *S_c_* and 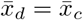, which means that the values for control can be directly compared to the values of untreated cell lines from other plates normalized by a standard Z-score. Visual comparison of raw and normalized data showed low variance in traces of the same compound while correctly preserving the relative changes of the compound traces to control (S5 Fig and S6 Fig).

Model output was also normalized to correspond to the normalized data. For untreated cell lines, we used Z-score as defined in Eq (5). For compound-treated cell lines, we used the adjusted Z-score whose form for the model output is

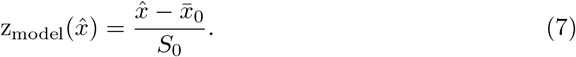

Here, 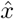 is model output, and 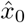 and *S*_0_ are sample mean and standard deviation of the simulation output for the control parameter set. Eq (7) can be derived directly from Eq (6). The model output is averaged over 1000 evaluations, which leads to low variance between repeated population-level simulations. Therefore, we set 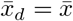, *S_d_* = *S_x_*, 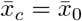, and *S_c_* = *S*_0_ which directly reduces Eq (6) to Eq (7).

### Optimization algorithm

The proposed mathematical model contains 6 free parameters: activator concentration (*A*), degradation rates (*d_M_*, *d_P_*, *d_R_*), light sensitivity (*i*) and noise intensity (*σ*). We adjusted those parameters to the untreated and compound-treated luminescence recordings in four consecutive steps using a differential evolution optimizer [49]. First, the noise was set to 0 and other parameters were estimated based on general properties of the circadian clock known from the literature. Second, we estimated noise based on the damping ratio observed in the luminescence recordings. Third, the model parameters were fine-tuned to the luminescence recordings of untreated cell lines. Fourth, we further adjusted the model parameters to the compound-treated cell lines. We developed this step-wise optimization approach to minimize the number of computationally expensive evaluations of the stochastic model, thus enabling the fitting of the stochastic model to a wide variety of experimental data. In the following, we describe the different steps of this fitting approach in detail.

#### Fitting the model in the absence of noise

In the first step of the fitting algorithm, a custom cost function was used to find a population of parameter sets that can reproduce basic properties of a circadian oscillator known from literature [50]. Those properties are oscillations with an approximately 24-hour period in constant darkness and entrainment by external LD cycles [20]. As the probability of the oscillations is the highest when the degradation terms are equal [51], we introduced a constrain on equal degradation terms

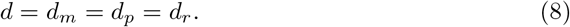

The cost function for optimization reads

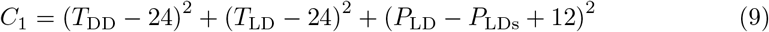

where *T*_DD_ is period in constant darkness (DD), *T*_LD_ is period under 12:12 LD cycle, *P*_LD_ is phase of the oscillator under LD, and *P*_LDs_ is phase of the oscillator under LD with 12-hour phase shift. First two terms of the equation ensure circadian oscillations in constant darkness and under 12:12 LD cycle respectively. The third term ensures that the model reacts to different phases of the LD cycles [50]. The optimization was repeated 50 times to obtain 50 different parameter sets. Those are consequently used as an initial population for the next fitting step.

#### Estimating noise intensity

Higher noise intensity causes faster desynchronization of individual oscillators, which leads to faster damping of oscillation amplitude in the population-level recordings [19]. We estimated the damping ratio from a damped sine fit to data. The function for the damped sine is

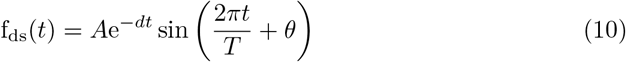

with amplitude *A*, damping ratio *d*, period *T*, and phase *θ*.

We estimated the corresponding value of the noise intensity parameter **σ** for each of the 50 parameter sets obtained in the previous fitting step. In this step, only the noise intensity was estimated while all other parameters of the model were fixed to the values estimated previously. The cost function for the differential evolution optimization is

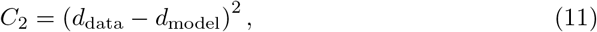

where *d*_data_ is the damping ratio estimated from the luminescence recordings (first six days of the fitting data, Fig 2A) and *d*_model_ is the damping ratio estimated from the model output (simulated in constant darkness after prior entrainment by 12:12 LD cycle).

#### Parameter adjustment for the untreated cell lines

To numerically find the optimal parameter set for a specific luminescence recording, we used differential evolution with the initial population given by the previous step of the optimization algorithm. As a cost function, we used the squared error

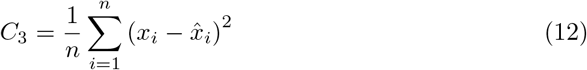

where *x_i_* denotes data points and 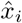 denotes the output of the model. The initial conditions were estimated by letting the model run under 12:12 LD cycles for 10 days. In this way, we diminished the effect of arbitrary initial conditions on the final solution. Further, we assumed that untreated cells produce sustained circadian oscillations in constant darkness [20]. Therefore, we checked the solution of the deterministic model (**σ** = 0) for the presence of sustained oscillations in constant darkness by detecting the amplitudes and locations of peaks. If no oscillations were found for the corresponding parameter set, the cost function returns its maximal value without the need to additionally evaluate the stochastic model with noise terms. Based on the assumption of sustained oscillations, we also kept the previously imposed constraint on the equality of degradation terms.

#### Parameter adjustment for the compound-treated cell lines

The cost function for experiments with pharmacological treatments is

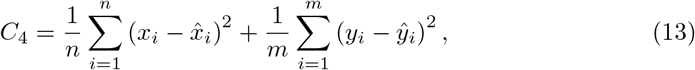

where *x_i_, y_i_* denote luminescence recordings recorded under the LD cycle and in constant darkness respectively and 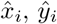 denote model outputs for the LD cycle and constant darkness respectively. For compound-treated cell lines, we could not assume that the cells exhibit oscillations in constant darkness. Therefore, we relaxed the constraint on equal degradation rates and did not test the deterministic model for the presence of sustained oscillations.

The compounds were administrated right before the beginning of the recording which caused transient behavior observable in the data. To incorporate this into our simulations, we precalculated initial conditions by running the model of the untreated cells for 10 days and then taking the last values as initial conditions for the simulations of pharmacological treatments. The resulted initial conditions correspond to the state of the oscillator at the transition from light to dark. This corresponds to the experimental data used in our work that starts at the beginning of the dark phase. The initial conditions were fixed for the whole process of optimization. Individual compound doses were fitted iteratively from control to the highest concentration of the specific compound. At each step, the final optimized population from the previous step was used as the initial population for the next step.

### Goodness of fit metric

We assessed the goodness of fit using the model efficiency coefficient defined as [52]

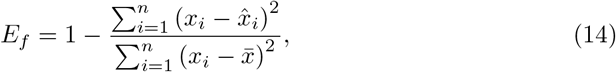

where *x_i_* are data points, 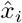 are corresponding model outputs and 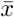 is data mean. The values of the model efficiency coefficient can range from −∞ to 1. Values near 1 indicate high predictive value of the model while negative values indicate that data mean 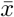 is a better predictor than model output 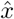.

### Software implementation

All code was written in Julia [53] version 1.5. For the numerical solution of the differential equations, we used the DifferentialEquations.jl package [54]. The fitting algorithm was implemented using differential evolution from the BlackBoxOptim.jl package. All data and code needed to replicate the results and generate the paper figures is available at https://github.com/vkumpost/biolum.

## Supporting information

**S1 Fig.**
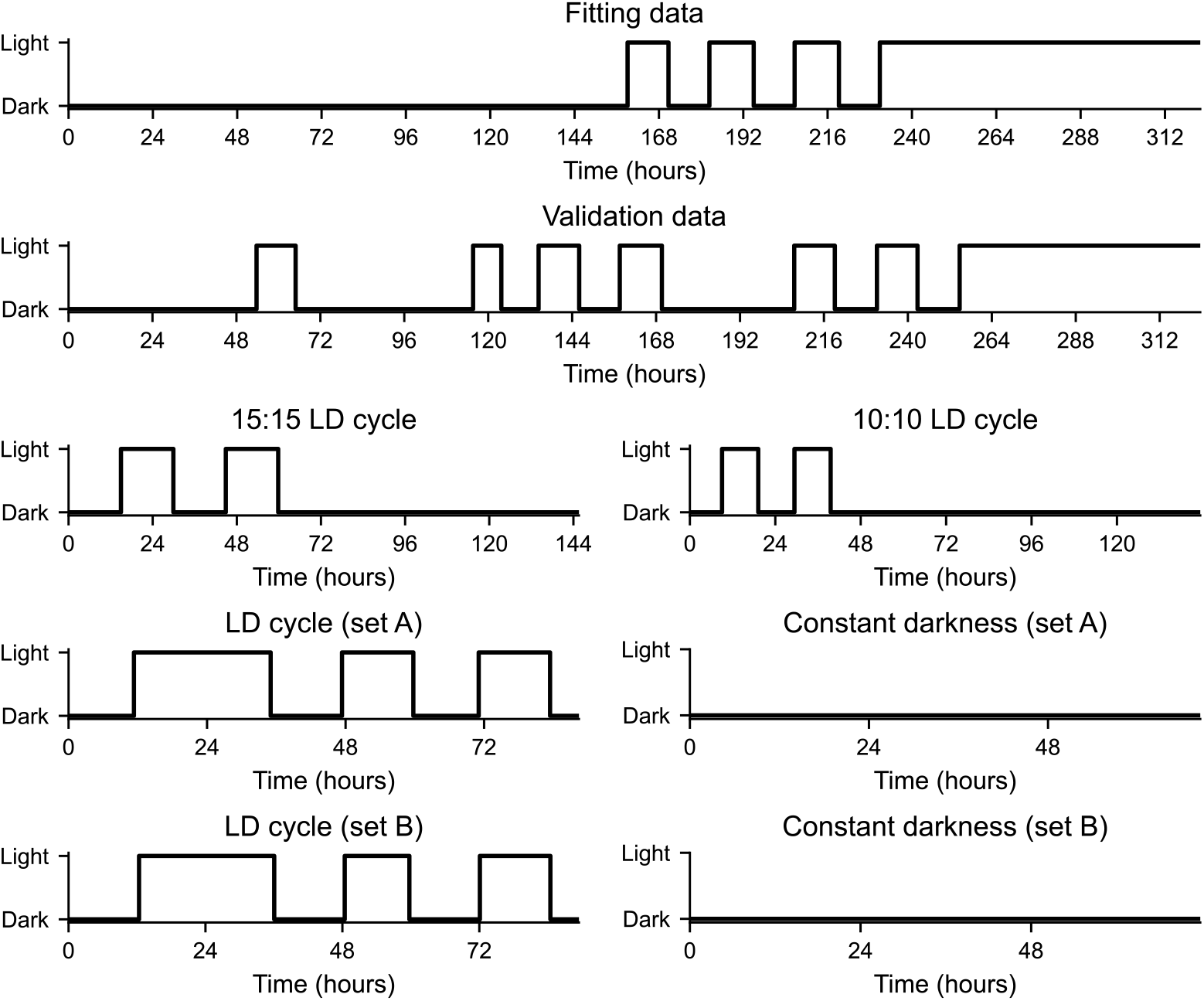
Lighting conditions. Fitting data was obtained by exposing untreated cell lines to constant darkness for six days, followed by three days of 12:12 LD cycle, and finally placed under constant light for three days. Validation data was recorded for 14 days under various lighting conditions, including LD cycles and periods of constant light and darkness. 15:15 LD cycle and 10:10 LD cycle data were recorded under two 15:15 and 10:10 LD cycles, respectively, followed by a period of constant darkness. Tested compounds were split into two plate sets (A and B). Each plate set consisted of two plates, one was placed under a light regime with a 24-hour phase reversal light pulse followed by two days of 12:12 LD cycle and the other was placed for three days in constant darkness.

**S2 Fig.**
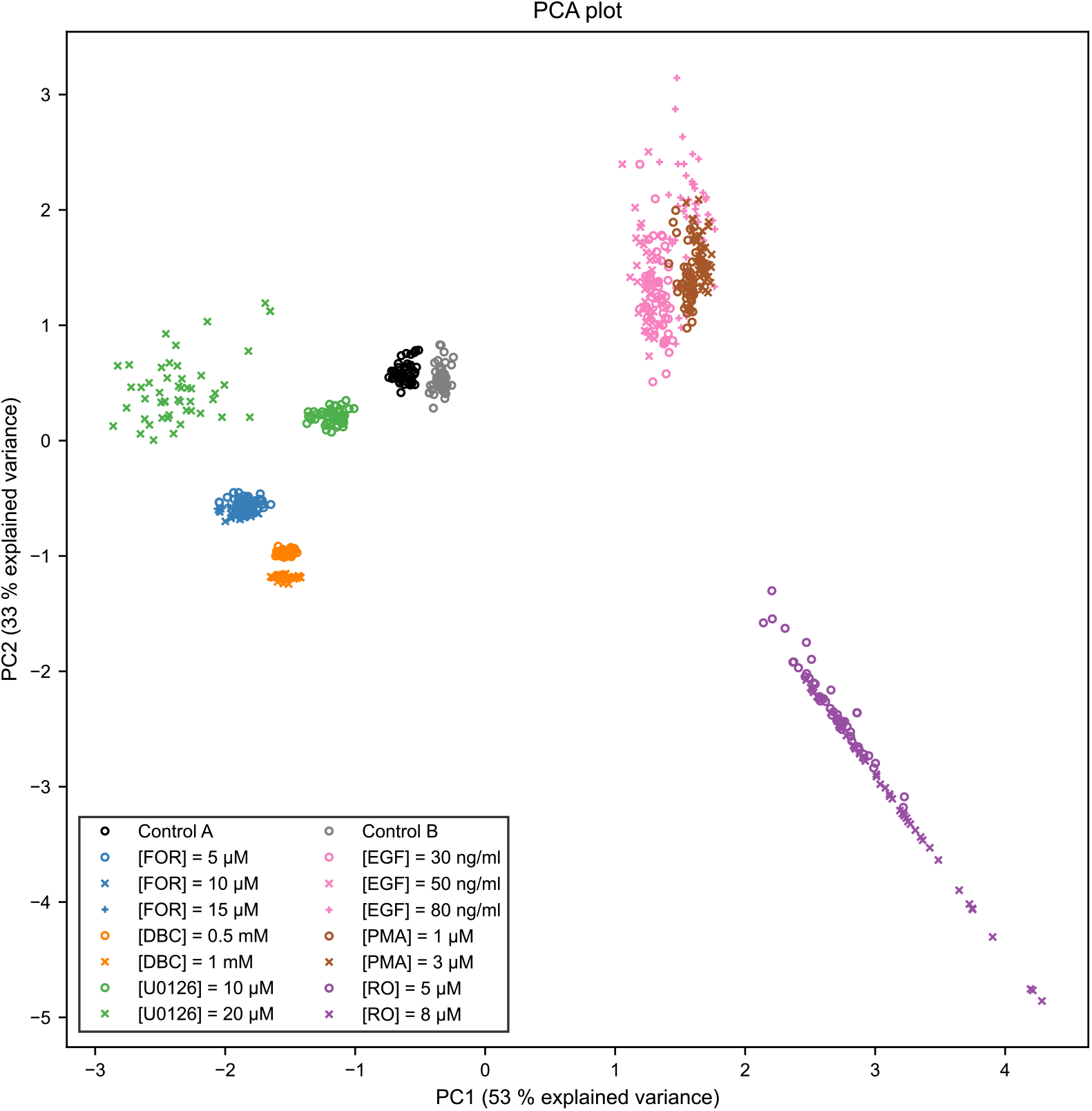
PCA plot – individual values. This plot is equivalent to Fig 3C showing individual points for each compound concentration. Each point represents one member of the final population obtained from running an evolutionary algorithm to find the best parameter fit to data. 50 agents were used as a fitting population for each compound.

**S3 Fig.**
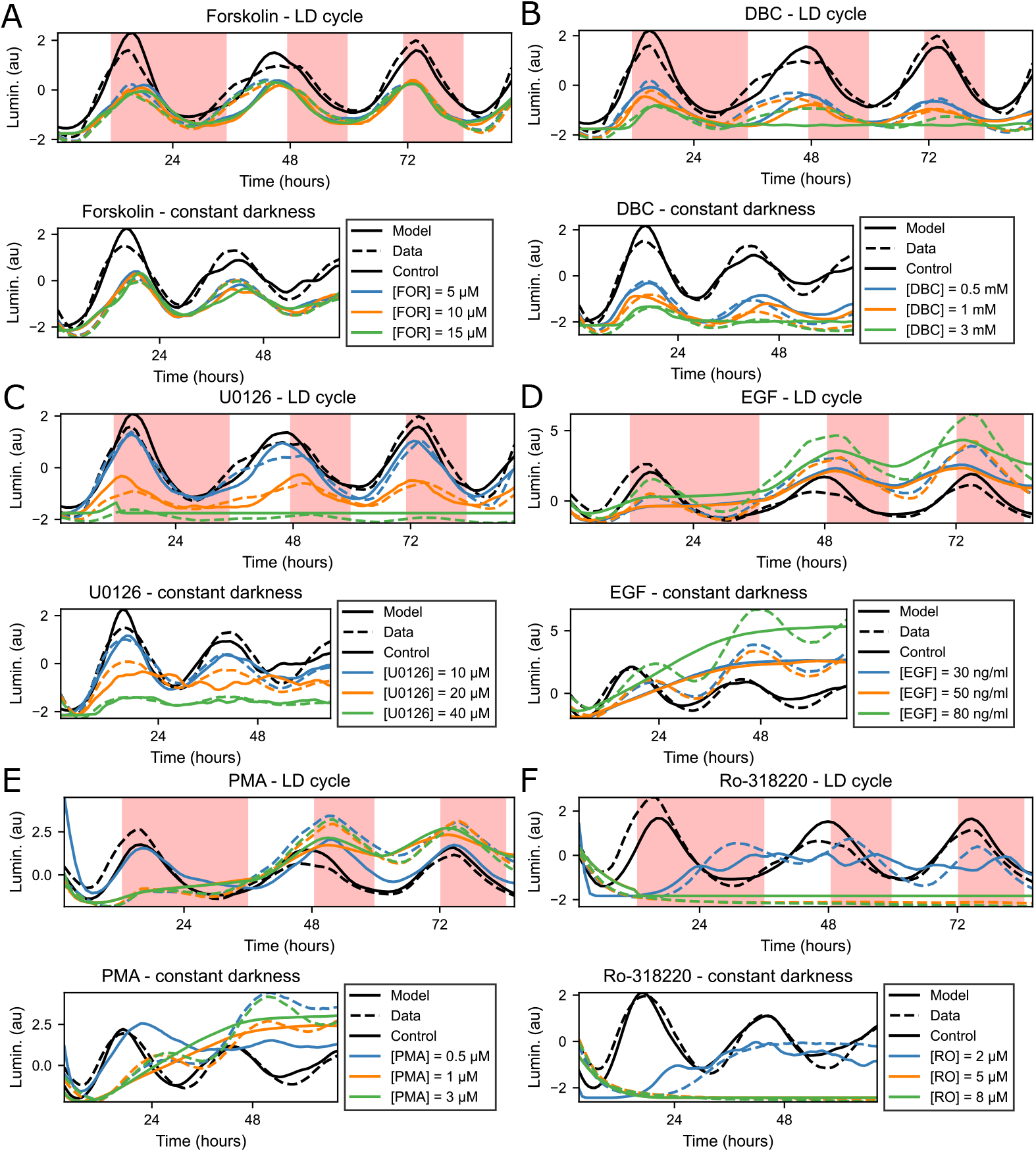
Model fit to pharmacological treatments. The model parameters were fitted to 6 compounds, each of which was applied in 3 different concentrations. The corresponding goodness of fit estimations can be seen in S2 Table. (A) Model fit to treatments with forskolin (FOR). (B) Model fit to treatments with dibutyryl cAMP (DBC). (C) Model fit to treatments with U0126. (D) Model fit to treatments with epidermal growth factor (EGF). (E) Model fit to treatments with phorbol-12-myristate-13-acetate (PMA). (F) Model fit to treatments with ro-318220 (RO).

**S4 Fig.**
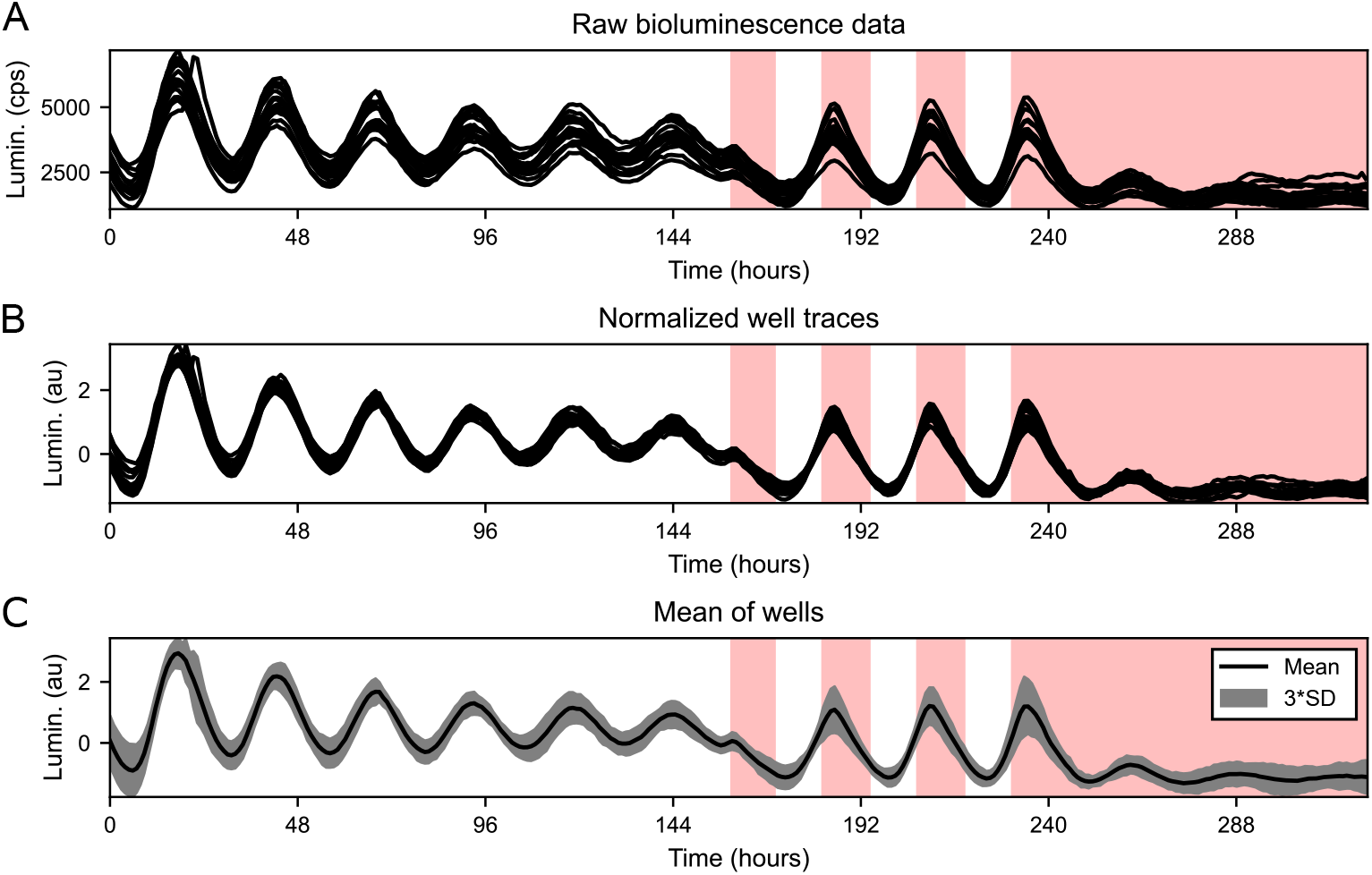
Normalization decreases variance in luminescence recordings. (A) Raw luminescence recordings in counts per seconds (cps) obtained from 16 cell culture wells. (B) After Z-score normalization, the variance among the individual traces is reduced and the resulting traces are presented in arbitrary units (au). (C) The individual normalized traces were used to calculate a mean value that we consequently used to fit and validate the model. The gray area around mean shows 3 * standard deviation (SD).

**S5 Fig.**
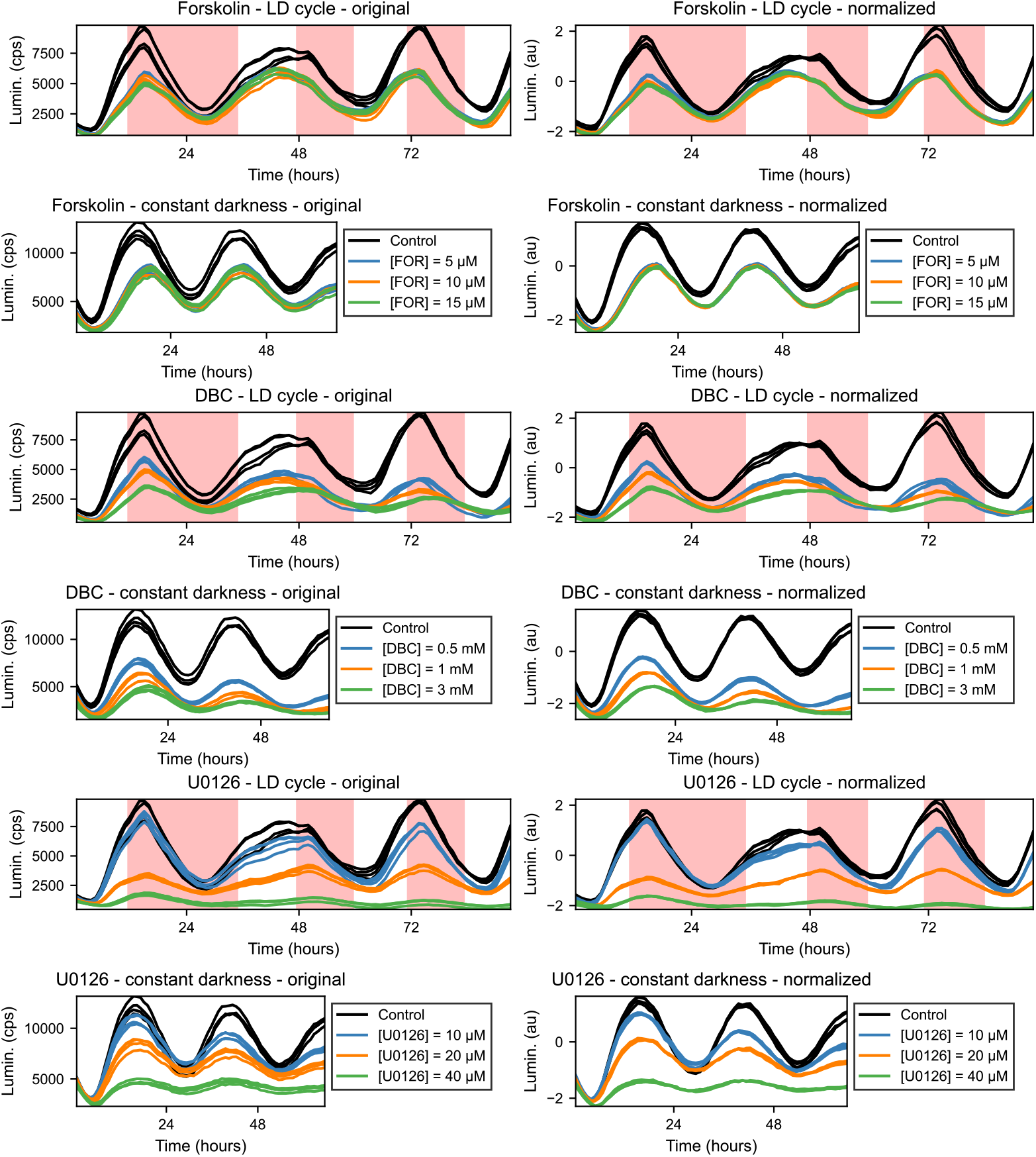
Original and normalized data for plate set A. The left column shows original unprocessed traces obtained from pharmacological treatments. In the right column are depicted the same traces after normalization by the adjusted Z-score. The unprocessed traces are shown in counts per second (cps) and the normalized traces are shown in arbitrary units (au).

**S6 Fig.**
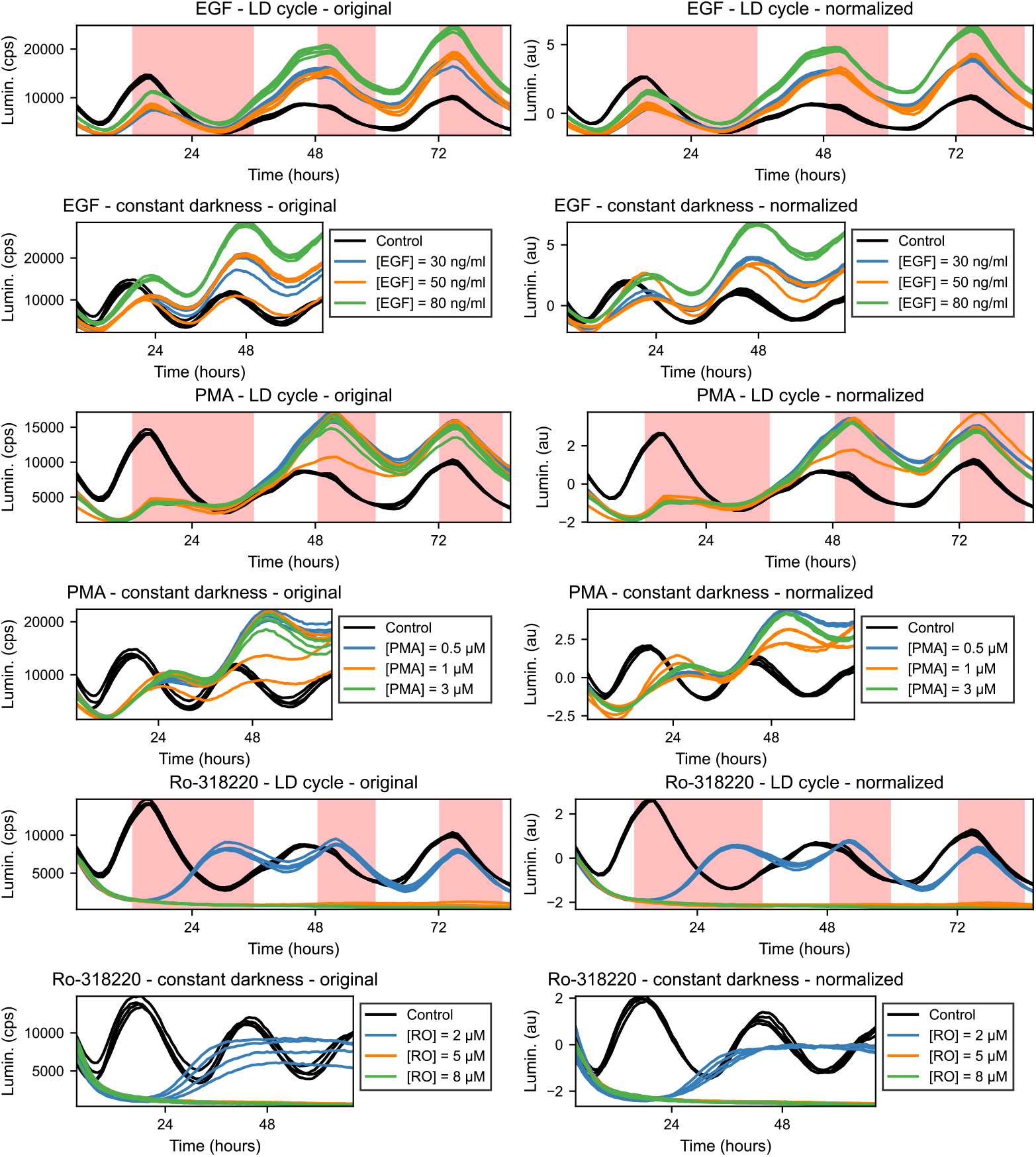
Original and normalized traces for plate set B. The left column shows original unprocessed traces obtained from pharmacological treatments. In the right column are depicted the same traces after normalization by the adjusted Z-score. The unprocessed traces are shown in counts per second (cps) and the normalized traces are shown in arbitrary units (au).

**S1 Table.** Well repeats. Description of well repeats on each plate used in our study. The pharmacological treatments were split into two plate sets (A, B).

| Plate | Well repeats |
| --- | --- |
| Fitting data | 16x untreated |
| Validation data | 4x untreated |
| 15:15 LD cycle | 32x untreated |
| 10:10 LD cycle | 16x untreated |
| LD cycle (set A) | 4x DMSO control<br>12x FOR (4x 5 μM, 4x 10 μM, 4x 15 μM)<br>12x DBC (4x 0.5 mM, 4x 1 mM, 4x 3 mM)<br>12x U0126 (4x 10 μM, 4x 20 μM, 4x 40 μM) |
| Constant darkness (set A) | 4x DMSO control<br>12x FOR (4x 5 μM, 4x 10 μM, 4x 15 μM)<br>12x DBC (4x 0.5 mM, 4x 1 mM, 4x 3 mM)<br>12x U0126 (4x 10 μM, 4x 20 μM, 4x 40 μM) |
| LD cycle (set B) | 4x DMSO control<br>12x EGF (4x 30 ng/ml, 4x 50 ng/ml, 4x 80 ng/ml)<br>12x PMA (4x 0.5 μM, 4x 1 μM, 4x 3 μM)<br>12x RO (4x 2 μM, 4x 5 μM, 4x 8 μM) |
| Constant darkness (set B) | 4x DMSO control<br>12x EGF (4x 30 ng/ml, 4x 50 ng/ml, 4x 80 ng/ml)<br>12x PMA (4x 0.5 μM, 4x 1 μM, 4x 3 μM)<br>12x RO (4x 2 μM, 4x 5 μM, 4x 8 μM) |

**S2 Table.**
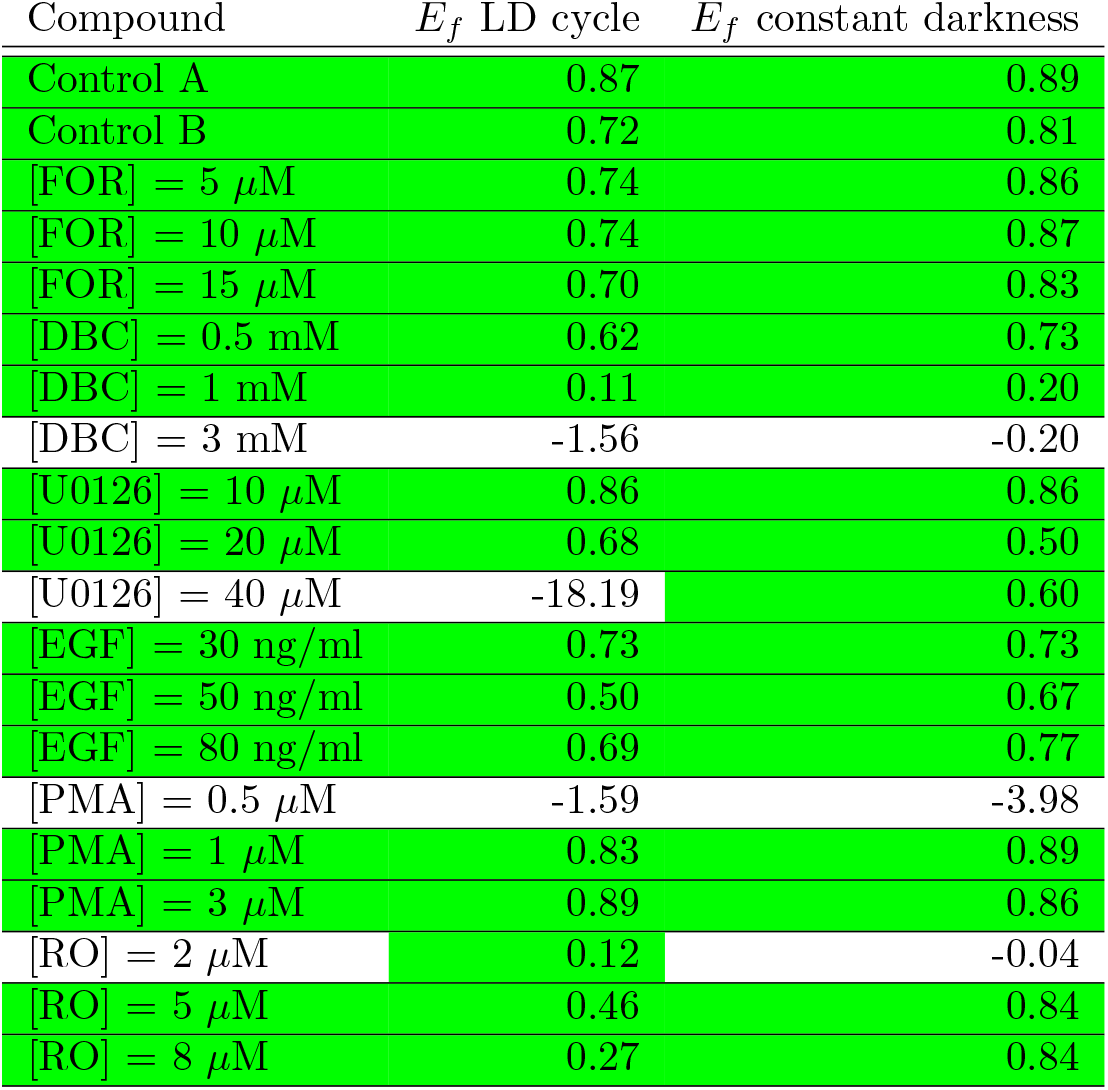
Goodness of fit for pharmacological treatments. The overall fit for a compound is considered good if *E_f_* for both LD cycle and constant darkness are positive (indicated by green color).

| Compound | $E_f$ LD cycle | $E_f$ constant darkness |
| --- | --- | --- |
| Control A | 0.87 | 0.89 |
| Control B | 0.72 | 0.81 |
| [FOR] = 5 μM | 0.74 | 0.86 |
| [FOR] = 10 μM | 0.74 | 0.87 |
| [FOR] = 15 μM | 0.70 | 0.83 |
| [DBC] = 0.5 mM | 0.62 | 0.73 |
| [DBC] = 1 mM | 0.11 | 0.20 |
| [DBC] = 3 mM | -1.56 | -0.20 |
| [U0126] = 10 μM | 0.86 | 0.86 |
| [U0126] = 20 μM | 0.68 | 0.50 |
| [U0126] = 40 μM | -18.19 | 0.60 |
| [EGF] = 30 ng/ml | 0.73 | 0.73 |
| [EGF] = 50 ng/ml | 0.50 | 0.67 |
| [EGF] = 80 ng/ml | 0.69 | 0.77 |
| [PMA] = 0.5 μM | -1.59 | -3.98 |
| [PMA] = 1 μM | 0.83 | 0.89 |
| [PMA] = 3 μM | 0.89 | 0.86 |
| [RO] = 2 μM | 0.12 | -0.04 |
| [RO] = 5 μM | 0.46 | 0.84 |
| [RO] = 8 μM | 0.27 | 0.84 |

## Acknowledgments

We would like to thank Irina Mamontova and Ines Reinartz for proofreading the manuscript. We are grateful for funding by the Helmholtz Association in the program Natural, Artificial and Cognitive Information Processing (VK, DV, NSF, RM, LH), HIDSS4Health - the Helmholtz Information & Data Science School for Health (VK, RM), and the Max Planck Society (DV, SBG, NSF). The funders had no role in study design, data collection and analysis, decision to publish, or preparation of the manuscript.

